# Mathematical modeling for a primitive form of habituation in an amoeba

**DOI:** 10.1101/2025.09.15.675989

**Authors:** Kota Nishi, Atsushi Tero, Yukinori Nishigami, Toshiyuki Nakagaki

## Abstract

Learning abilities, once thought to be unique to higher animals, have been reported to exist in their primitive form in single-celled organisms. This has triggered a growing interest in carefully examining the nature and mechanisms of the primitive versions of learning abilities, which would provide important clues for understanding the evolution of behavioral capabilities in organisms. In this study, we focused on previous experimental studies showing that the slime mold *Physarum polycephalum*, a model organism for studying protist behavior, exhibits the ability to adapt to chemical environments. We propose a possible dynamic mechanism underlying this habituation, reproducing reported experimental observations with accuracy. By refining a mathematical model that was as simple as possible and based on non-specific biochemical processes within cells, we clarified a plausible mechanistic framework. Based on these results, we examined the similarities and differences between this framework and previously proposed habituation models of single-cell movement and animal neural-circuit regulation. These findings are significant because they open new avenues for research into the generality and evolutionary origins of acclimation learning.

## 1 Introduction

Studies on information processing in simpler organisms, such as single-celled protists that can evoke an adaptive response to complex environmental situations, have deepened our understanding of cognitive behavior in neural organisms (Lyon et al. 2021). Several types of adaptive behavior have been reported in the amoeboid organism *Physarum polycephalum*, making it an important model organism for elucidating a primitive form of cognitive function in higher animals (Reid 2023).

Plasmodium of *P. polycephalum* is an amoeba-like multinucleated unicellular organism with a large sheet-like body up to several m^2^. Its protoplasm exists in two phases; protoplasmic gel (ectoplasm), which exhibits passive elasticity and active rhythmic contractility, and protoplasmic sol (endoplasm), which is a viscous liquid. Inside the sheet-like body of plasmodium, an intricate network of veins (tubular channels for protoplasmic streaming) develops toward the rear from the extending front.

The ectoplasmic gel periodically contracts based on the biochemical regulation of the actomyosin system, and these self-sustained oscillations of contraction show some phase and force differences within the body. As this contraction generates pressure on the sol, these differences cause it to flow within the body through the tube network. This sol flow causes amoeboid migration of plasmodium while inducing major morphological changes in the tube network.

Plasmodium can solve difficult foraging problems by reforming its efficient tube-network (Nakagaki et al. 2000; Tero et al. 2010), and can memorize and anticipate the environmental periodic changes as well (Saigusa et al. 2008). These adaptive behaviors and their underlying mechanisms have been analyzed using mathematical modeling (Nakagaki et al. 2007; Saigusa et al. 2008; Tero et al. 2007, 2010).

As this mathematical model elucidated the key mechanism underlying adaptive behavior, it was applicable to similar types of behavior in higher animals, such as the social design of public transportation networks in human society (Tero et al. 2010) and ant trail formation in complicated landscapes (Ma et al. 2013).

As discussed above, studies on *P. polycephalum* have provided a new perspective on the evolution of behavioral intelligence (right from cells to animal societies). Along this line of study, we aimed to investigate habituation in *P. polycephalum* and analyze the underlying mechanism by mathematical modeling.

Habituation, a type of learning, refers to the decrease in behavioral responses when stimuli are repeatedly presented (Rankin et al. 2009). Habituation is distinct from sensory adaptation, sensory fatigue, and motor fatigue (Rankin et al. 2009). Several studies have attempted to elucidate the mechanisms underlying habituation. Most habituation studies have focused on multicellular organisms that possess a nervous system (Castellucci et al. 1970; Johnson and Wuensch 1994).

However, some single-celled microorganisms, such as ciliates (Wood 1988a,b; Rajan et al. 2023) and amoebae (Boisseau et al. 2016), also exhibit habituation. Habituation to mechanical stimuli has been extensively demonstrated in ciliates (Dussutour 2021). For example, in the ciliate *Stentor*, the contraction response to mechanical stimulation is an example of habituation (Wood 1988a). In this habituation, changes in intracellular Ca^2+^ concentration are crucial (Dussutour 2021).

In amoebae, habituation to chemical stimulation has been observed in plasmodium of *P. polycephalum*, as demonstrated by the hallmarks of habituation (Boisseau et al. 2016). However, the underlying mechanisms have not been adequately explored. Therefore, in this study, we propose a mathematical model to reproduce habituation in *P. polycephalum*. We compared this model with previous models and discussed the commonalities in the habituation mechanism.

## 2 Nature of Habituation in Plasmodium and Aim of Mathematical Modeling

Plasmodium usually locomotes when searching for food products. It decreases its speed when encountering quinine, a chemical repellent (Takagi et al. 2007). Boisseau et al. (2016) demonstrated the hallmarks of habituation in plasmodium by leveraging this property. They conducted bridge-crossing experiments on plasmodium (Fig. 1). In their experiment, a plasmodium was placed on a food patch, with a new food patch connected by one of two agar bridges: an agar bridge containing quinine (hereinafter, b-Q) or a normal agar bridge without quinine (hereinafter, b-N). The bridge was set between plasmodium and the new food patch, and plasmodium was then made to cross either the b-Q or the b-N once per day (Fig. 1). They recorded the time from when the tip of plasmodium entered the bridge to when it reached a new food patch (hereinafter called the time to cross the bridge). From their experiments, the following observations were made: (i), the time taken to cross the b-Q decreased when plasmodium repeatedly crossed the b-Q (Fig. 2(a)), (ii), the time to cross the b-Q recovered after plasmodium crossed the b-N for 2 consecutive days (Fig. 2(a)), (iii), the time required to cross the b-N does not depend on the number of previous passes through the b-Q (Fig. 2 at day 7). In particular, features (i) and (ii) correspond to habituation and spontaneous recovery, the hallmarks of habituation (Table 1), respectively. Although feature (iii) is not directly related to habituation, feature (iii) suggests that plasmodium reflects the stimulus history of its motility only under necessary circumstances.

**Table 1.** The hallmarks of habituation (excerpt)

| Name | Feature |
| --- | --- |
| habituation | Repeated application of a stimulus results in a progressive decrease in some parameter of a response to an asymptotic level. |
| spontaneous recovery | If the stimulus is withheld after response decrement, the response recovers at least partially over the observation time. |

**Fig. 1.**
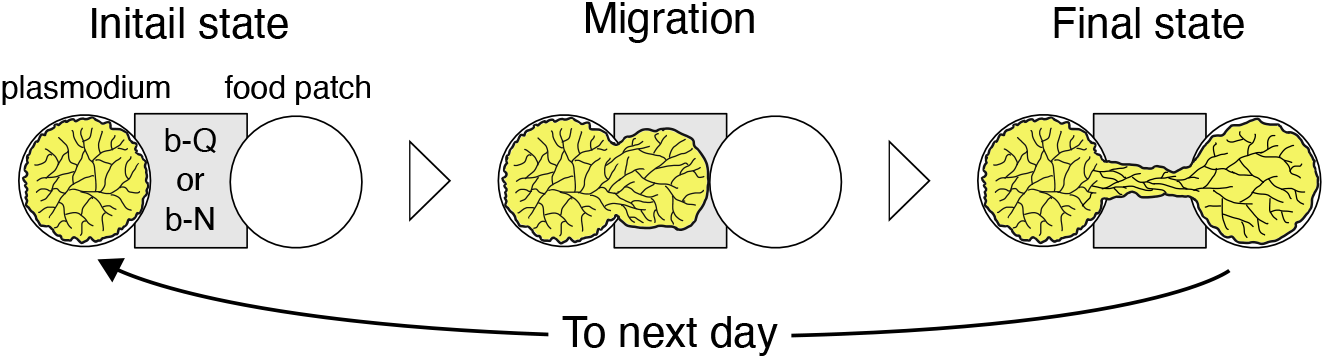
Illustration depicting the workflow of the bridge crossing experiment conducted by Boisseau et al. (2016). The b-Q represents the agar bridge containing quinine, and the b-N represents the normal agar bridge

**Fig. 2.**
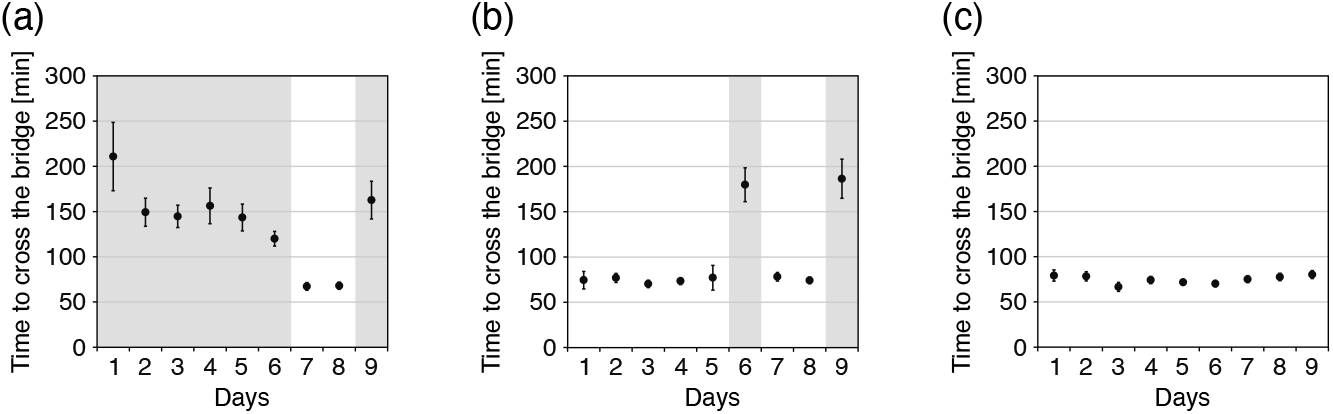
Time taken by plasmodium to cross the bridge (from day 1 to 9) (Boisseau et al. 2016). Black point represents the mean time to cross the bridge, and the error bar represents the 95 percent confidence interval: the gray background represents the day when plasmodium crossed the b-Q, and the white background represents the day when plasmodium crossed the b-N. Data adapted from Boisseau et al. (2016)

This table adapted from Rankin et al. (2009).

In this study, we developed a mathematical model to account for the three behavioral features (i) ∼ (iii). We formulated a locomotion model of plasmodium driven by a chemical reaction–diffusion process. In the next section, we describe how the physiological and behavioral properties of plasmodium are represented by a mathematical model.

## 3 Formulation of the Mathematical Model

### 3.1 Overview of Our Model

First, we present an overview of the formulated model. We developed the following model, which comprises the equation describing plasmodium locomotion (Eq. (1)) and the reaction–diffusion system regulating locomotion (Eq. (2) and (3)).

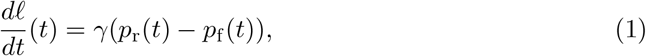

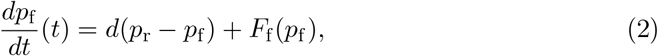

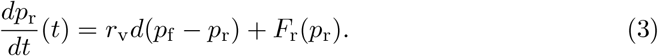

where

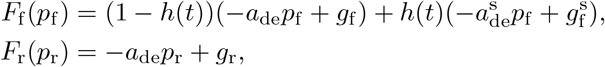

*ℓ* represents the position of plasmodium tip and *p*_f_ and *p*_r_ represent the sol pressures at the front and rear parts, respectively (Fig. 3). Function *h*(*t*) equals 1 in the presence of stimulation and 0 in its absence. Plasmodium expands its tip through sol flow in the tube network. The sol flow is driven by the gradient of the sol pressure. Therefore, we assumed that the tip expansion velocity is proportional to the pressure gradient (Eq. (1)). In this study, plasmodium locomotion was restricted to a straight line. In addition, we regarded plasmodium as consisting of two parts: the front part, which is the extendable tip for locomotion, and the rear part, which is the section opposite to the tip (Fig. 3). Accordingly, we introduced the sol pressure variable at each part, *p*_f_ and *p*_r_. Thus, *p*_r_ − *p*_f_ represents the pressure gradient (Eq. (1)). The locomotion model (Eq. (1)) indicated that plasmodium migration is driven by the difference between the pushing force from the rear, which promotes forward movement, and the pushing-back force from the front, which inhibits it. By identifying the sol pressure with the concentration of the chemical involved, we formulated the dynamics of the sol pressure using a simplified reaction–diffusion system (Eq. (2) and (3)). The first term on the right-hand side is the diffusion term and the second term is the reaction term (Eq. (2) and (3)). Here, the parameters of the reaction term at the front switch in the presence or absence of stimulation because plasmodium detects the repellent at the front.

**Fig. 3.**
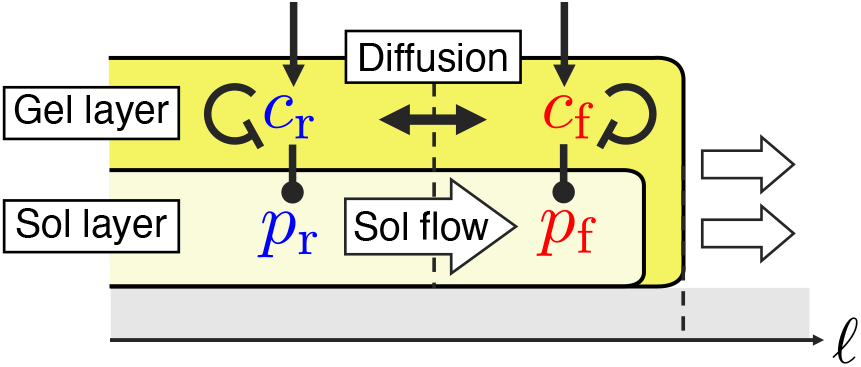
Schematic illustration of a cross-section of plasmodium. Each of *c*_f_ and *c*_r_ represents a chemical that regulates the gel contraction. *p*_f_ and *p*_r_ represent the sol pressure. *ω* represents the coordinate of the tip position. A line with a circle represents an activation effect, a line with a T-shaped end represents an inhibition effect, an arrow represents the generation effect of a chemical, and a double-headed arrow represents the diffusion effect

We developed a plasmodium locomotion model driven by a special difference in the chemical concentration that regulates the sol pressure. The stimulation directly influenced the chemical reaction at the front, whereas the reaction at the rear was indirectly affected by the diffusion between the front and rear. In the following section, we describe the model formation in more detail.

### 3.2 Detail of Formulation

First, we derived a locomotion model in plasmodium. We assumed that the tip expansion velocity was proportional to the volumetric flow rate of the sol toward its tip because plasmodium expanded its tip due to the sol flow in the tube network. Thus, we formulated the tip expansion velocity (hereafter called the locomotion velocity) as follows:

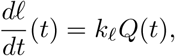

where *Q*(*t*) denotes the volumetric flow rate of the sol. The sol flow inside the tube can be approximated using a Hagen-Poiseuille flow: The volumetric flow rate is expressed as follows:

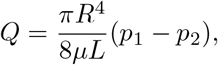

where *Q* is the volumetric flow rate, *R* is the tube radius. *L* is the tube length, *µ* is the viscosity coefficient, and Here, *p*_1_ and *p*_2_ denote the pressures at the tube edges. In this study, we considered plasmodium to be composed of two parts: front and rear (Fig. 3). We assumed that *n* tubes cross between the front and rear parts and that they are identical. Therefore, the volumetric flow rate of the sol toward the front part is as follows:

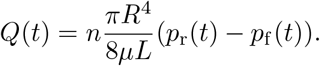

Consequently, the locomotion velocity of plasmodium is represented as

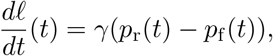

where

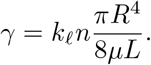

Next, we formulated a reaction–diffusion model for chemically involved gel contraction with simplified diffusion terms. We considered the chemical reactions, generation and decomposition, and the chemical diffusion between the front and rear parts. Then, the reaction–diffusion model was formulated as follows:

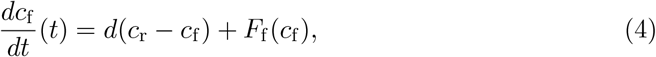

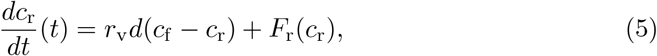

where

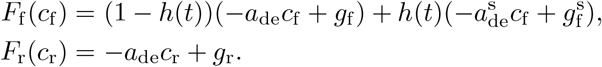

The variables *c*_f_ and *c*_r_ represent the concentrations of a chemical involved in gel contraction in the front and rear parts, respectively. The terms *d*(*c*_r_ − *c*_f_) and *r*_v_*d*(*c*_f_ − *c*_r_) represent the chemical diffusion between the front and rear parts, respectively, where *r*_v_ is their volume ratio. Note that we assume diffusion between solutions with different volumes, and therefore diffusion terms describe above (Appendix A). The terms *F*_f_ (*c*_f_) and *F*_r_(*c*_r_) represent chemical reactions. 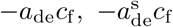 and −*a*_de_*c*_r_ represent the decomposition of chemicals and 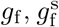 and *g*_r_ represent the generation of the chemicals. Function *h*(*t*) equals 1 in the presence of stimulation and 0 in its absence. Thus, the chemical reaction at the front switches depending on its presence or absence.

Finally, we derived the temporal dynamics of the sol pressure. Sol pressure was generated by gel contraction. In this study, we assumed a linear relationship between sol pressure and the concentration of a chemical involved in gel contraction, although a more complex relationship may exist. From this assumption, the following equations were obtained:

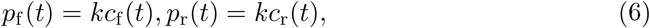

where *k* denotes a positive constant. From Eq. (4), (5), and (6), the sol-pressure dynamics were derived as follows:

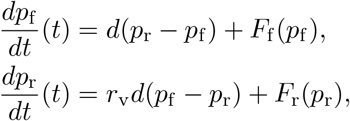

where

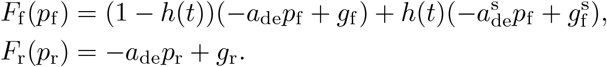

Here, 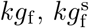 and *kg*_r_ replace 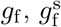 and *g*_r_, respectively.

In conclusion, the model presented in Sec. 5.1. In the next section, we describe simulations performed using this model.

## 4 Simulation

First, we explain the simulation settings using a simple case study (Fig. 4). The purpose of our simulation was to compute the time required to cross the bridge daily, as in Boisseau et al. (2016). First, given the initial value of (*p*_f_ (*t*), *p*_r_(*t*)) for each day, we compute the crossing time using Eq. (1), (2), and (3) (Fig. 4). We applied stimulation in the following pattern, depending on the bridge type. If the bridge type is the b-Q on day *n* (*t*_*n*−1_ ≤ *t < t*_*n*_),

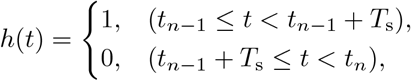

where *t*_*n*_ = 24*n*, 0 *< T*_s_ *<* 24. If the bridge is the b-N on day *n*,

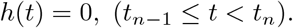

**Fig. 4.**
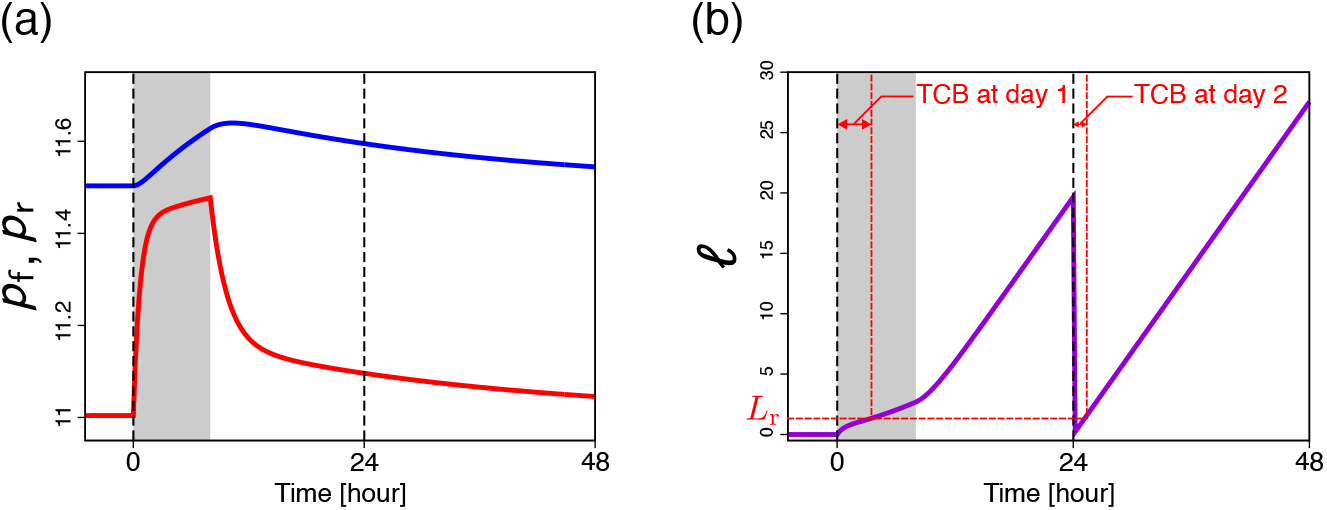
Time course of our model (Eq. (1), (2), and (3)) where the bridge condition at day 1 is the b-Q, and that at day 2 is the b-N. (a) time course of sol pressure. The red line represents *p*_f_, and the blue line represents *p*_r_. (b) time course of the tip position of plasmodium. The value of *ω* resets to 0 at the start of the day. *L*_r_ represents the bridge length. TCB means the time to cross the bridge. Gray background represents the term under stimulation, *h*(*t*) = 1, and white background represents the term in the absence of stimulation, *h*(*t*) = 0

The initial value of (*p*_f_ (*t*), *p*_r_(*t*)) on Day 1 is 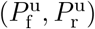, which is the equilibrium state in the absence of stimulation. After Day 2, the final states of (*p*_f_ (*t*), *p*_r_(*t*)) on the previous day were used as the initial conditions (Fig. 4(a)). Second, we computed the time taken to cross the bridge as the time required for *ℓ* (*t*), the position of plasmodium tips, to reach *L*_r_, the length of the bridge, on each day (Fig. 4(b)).

Our simulation results reproduced three features (i) ∼ (iii) observed in the experimental results (Fig. 5). We focused on the temporal dynamics of the tip expansion velocity, corresponding to Fig. 5(a) (Fig. 6). Mitigation of deceleration due to repeated stimulation, followed by recovery of deceleration after a temporary interruption in stimulation, was observed (Fig. 6). In addition, the velocity immediately returned to 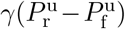 when the stimulation was withheld (Fig. 6). The mitigation of deceleration under stimulation indicated a decline in the time to cross the b-Q, and its recovery indicated an increase in the time to cross the b-Q (Fig. 5(a)). In addition, the immediate return of velocity to 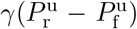 corresponds to the consistency of the time required to cross the b-N (Fig. 5). In other words, these temporal behaviors enable the reproduction of three features (i) ∼ (iii). Therefore, in the subsequent section, we discuss the relationship between the sol pressures (*p*_f_, *p*_r_) and the velocity dynamics.

**Fig. 5.**
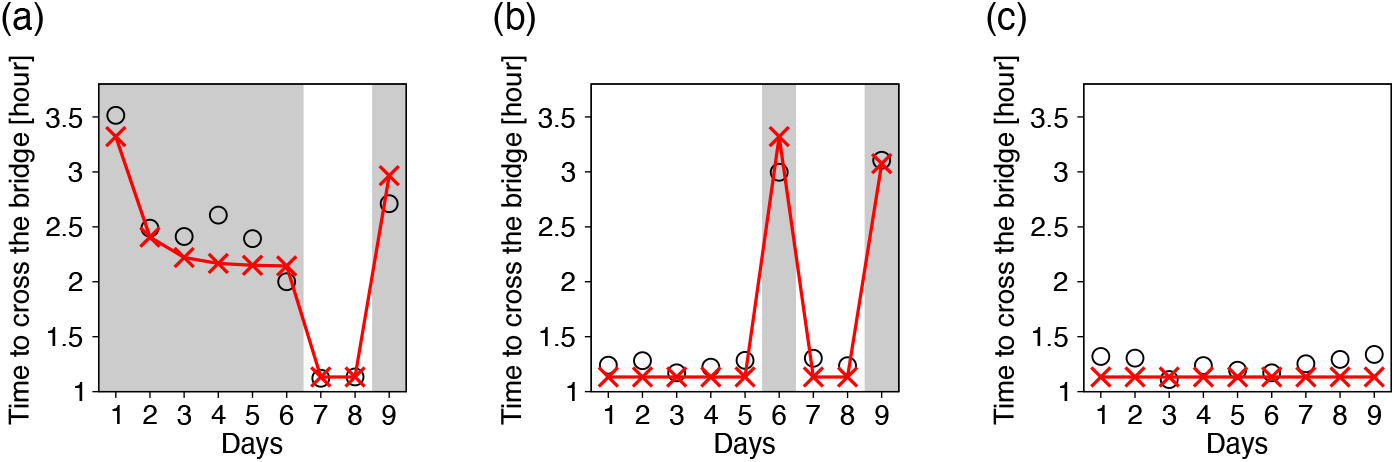
Our simulation results of time to cross the bridge. The Red cross mark represents our simulation results, and the circle represents the mean time to cross the bridge in the experiment (Boisseau et al. 2016). Gray background represents the day when the bridge condition is quinine, and white background represents the day when the bridge condition is normal. (Parameters: 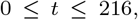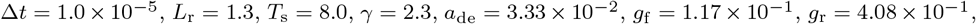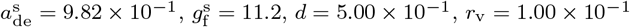)

**Fig. 6.**
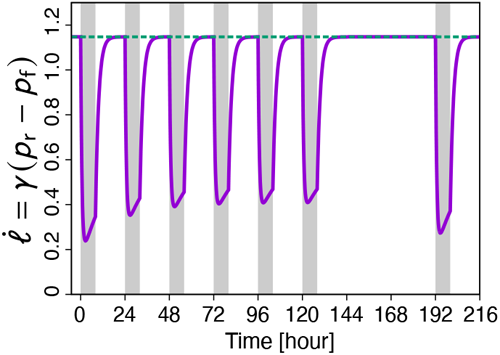
The temporal dynamics of the tip expansion velocity corresponding to Fig. 5(a). Gray background represents the term under stimulation, *h*(*t*) = 1, white background represents the term in the absence of stimulation, *h*(*t*) = 0, and green broken line represents that the velocity is equal to 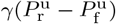

## 5 Mathematical Analysis

We analyzed the behavior of the locomotion velocity from the behavior of the sol pressure on the (*p*_f_, *p*_r_) plane. At this point, the dynamics of the sol pressure in the presence and absence of stimulation on the (*p*_f_, *p*_r_) plane and the relationship between the pressure and locomotion output are required. In Sec. 5.1, we analyze the dynamics of the pressure on the (*p*_f_, *p*_r_) plane and the relationship between the pressure and locomotion output. In Sec. 5.2 and 5.3, we discuss the behavior of the locomotion velocity based on the analysis in Sec. 5.1.

### 5.1 Mathematical Preparation

First, we analyzed the dynamics of the sol pressure. In the presence of a stimulation (*h*(*t*) = 1), Eq. (2) and (3) can be transformed as follows.

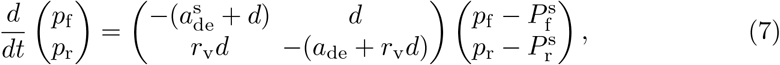

where 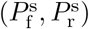 denotes the equilibrium point.

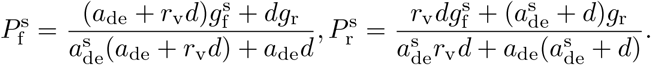

The eigenvalues and eigenvectors in Eq. (7) are

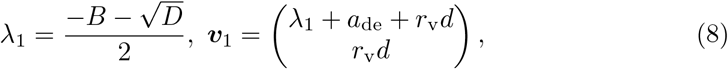

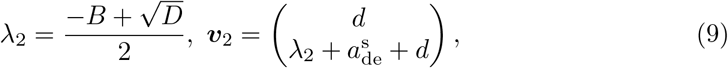

where *λ*_*i*_ represents the eigenvalue, ***v***_*i*_ represents the eigenvector corresponding to eigenvalue *λ*_*i*_, and

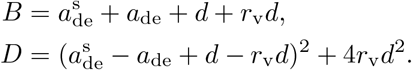

The dynamics of Eq. (7) depict in Fig. 7(a). In the absence of a stimulation (*h*(*t*) = 0), Eq. (2) and (3) can be modified as follows:

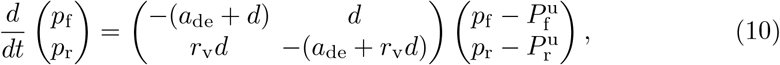

where 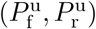 denotes the equilibrium point.

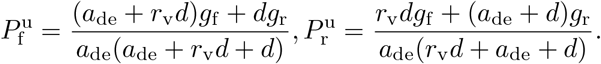

**Fig. 7.**
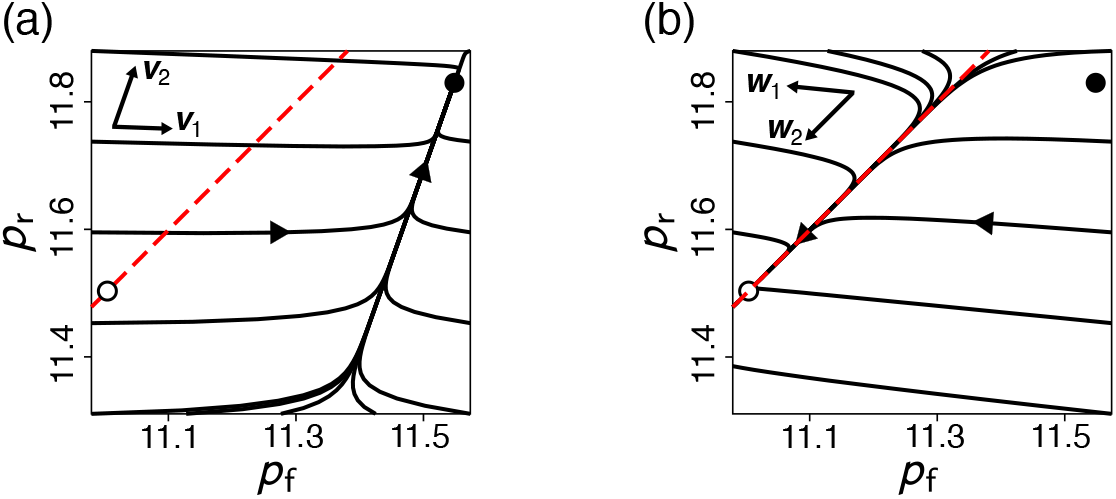
(a) The solution trajectory of Eq. (7). The arrows at top-left represent the directions of eigenvector ***v***_1_ and ***v***_2_ (Eq. (8) and (9)). (b) The solution trajectory of Eq. (10). The arrows at top-left represent the directions of eigenvector ***w***_1_ and ***w***_2_ (Eq. (11) and (12)). The red broken line represents the straight line 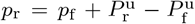. The white circle represents the equilibrium point in the absence of stimulation, 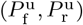, and the black circle represents the equilibrium point under stimulation, 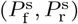. The parameters are same at Fig. 5

The eigenvalues and eigenvectors in Eq. (10) are

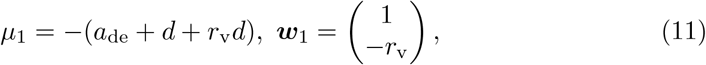

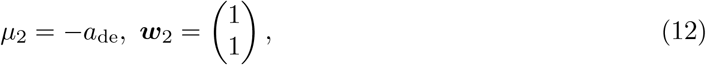

where *µ*_*i*_ represents the eigenvalue, and ***w***_*i*_ represents the eigenvector corresponding to the eigenvalue *µ*_*i*_. The dynamics of Eq. (10) depict in Fig. 7(b).

Second, we analyzed the relationship between the sol pressure (*p*_f_, *p*_r_) and behavioral output on the (*p*_f_, *p*_r_) plane. From Eq. (1), we define the function *V* (*p*_f_, *p*_r_) that returns the locomotion velocity corresponding to the sol pressure (*p*_f_, *p*_r_).

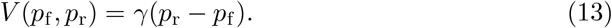

From Eq. (13), *V* (*p*_f_, *p*_r_) has a constant value on the line

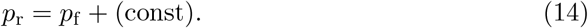

For example, *V* (*p*_f_, *p*_r_) = 1 for all (*p*_f_, *p*_r_) satisfying *p*_r_ = *p*_f_ +1*/ε*. The lines in Eq. (14) is referred to as the iso-velocity line. The iso-velocity lines are shown in Fig. 8(a). Moreover, by differentiating both sides of Eq. (1), and substituting Eq. (7), we obtain the acceleration under stimulation as follows:

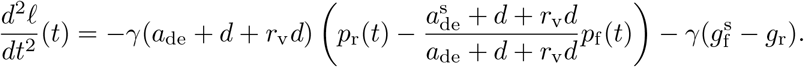

**Fig. 8.**
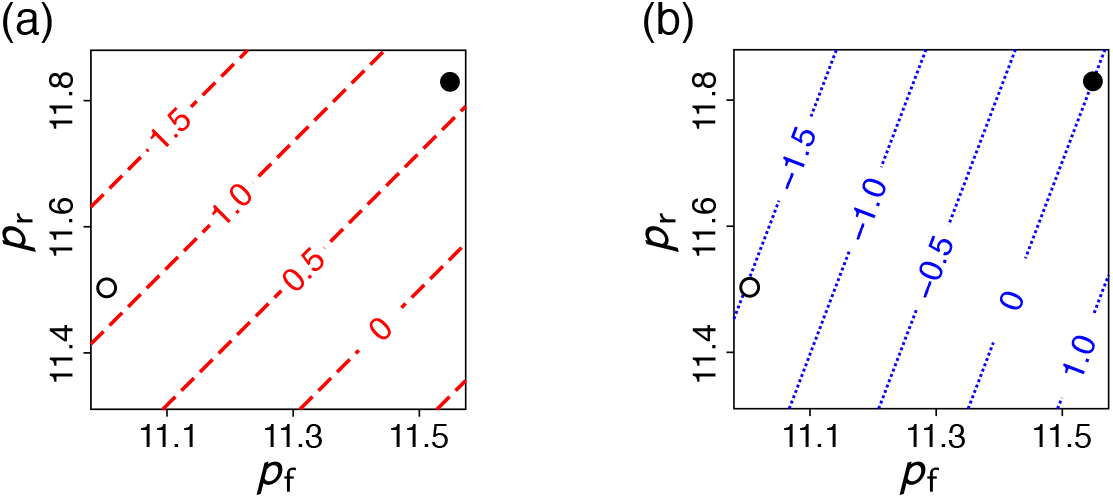
(a) The plot of *V* (*p*_f_, *p*_r_) on the (*p*_f_, *p*_r_) plane. Red broken lines represent the iso-velocity lines. The value on each line represents the velocity corresponding to the sol pressure (*p*_f_, *p*_r_) on the line. (b) The plot of *A*_s_(*p*_f_, *p*_r_) on the (*p*_f_, *p*_r_) plane. Blue dotted lines represent the iso-acceleration lines. The value on each line represents the acceleration under stimulation corresponding to the sol pressure (*p*_f_, *p*_r_) on the line. The white circle represents 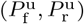, and the black circle represents 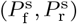. The parameters are same at Fig. 5

We define the function *A*_s_(*p*_f_, *p*_r_), which returns the acceleration under stimulation for sol pressure (*p*_f_, *p*_r_).

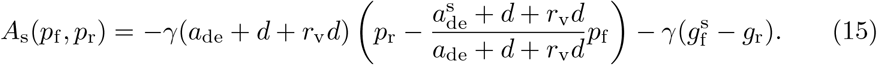

From Eq. (15), *A*_s_(*p*_f_, *p*_r_) has a constant value on the line

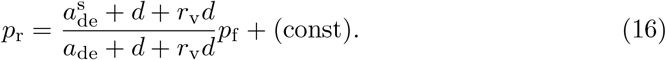

The lines in Eq. (16) is referred to as the iso-acceleration line. The iso-acceleration lines are shown in Fig. 8(b). Because 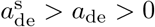 holds for our simulation parameters, the slope of the iso-acceleration line is greater than 1 (Eq. (16)). Hence, the iso-acceleration and iso-velocity lines intersect, as shown in Fig. 9.

**Fig. 9.**
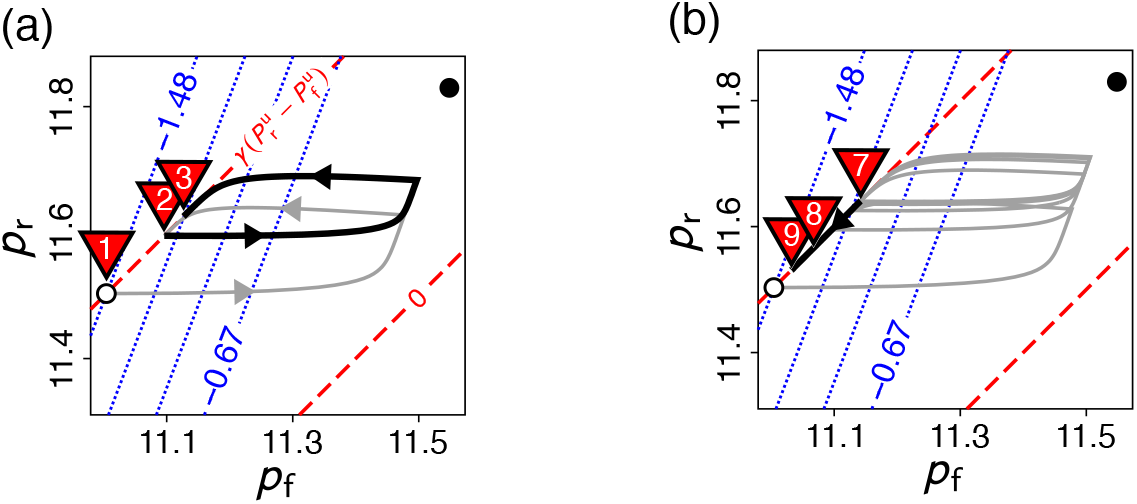
The sol pressure dynamics on the (*p*_f_, *p*_r_) plane corresponding to Fig. 5(a). (a) The trajectory of day 1*∼*2: the gray solid line represents the trajectory of day 1 (b-Q), and the black solid line represents the trajectory of day 2 (b-Q). (b) The trajectory of day 1*∼*8: the gray solid line represents the trajectory of day 1*∼*6 (b-Q), and the black solid line represents the trajectory of day 7*∼*8 (b-N). The white circle represents 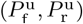, the black circle represents 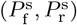, red broken lines represent the iso-velocity lines, blue dotted lines represent the iso-acceleration lines, and the red triangle with integer *n* points to the initial state at day *n*

### 5.2 Mitigation and Recovery of Deceleration

In this section, we discuss the mitigation and recovery of the deceleration under stimulation. The shift in the initial pressure state each day was crucial for the mitigation and recovery of deceleration under stimulation. By repeatedly crossing the b-Q, the initial state of each day shifts to the upper right (Fig. 9(a)). Focusing on the iso-acceleration lines, the initial state transitions in the direction in which the acceleration increases (Fig. 9(a)). It follows that the deceleration is mitigated (Fig. 6). However, the initial state shifted to the lower left each day when crossing the b-N (Fig. 9(b)). In other words, the initial state transitions in the opposite direction when repeatedly crossing the b-Q. Thus, deceleration under stimulation recovered when crossing the b-N. In summary, the mitigation of deceleration when repeatedly crossing the b-Q corresponded to an upper right shift of the initial state (Fig. 9(a)), and its recovery when crossing the b-N corresponded to a lower left shift (Fig. 9(b)).

Next, we discuss why the initial state shift occurs when the crossing the b-Q is repeated. To focus on the displacement in the direction along the iso-velocity line, namely, (1, 1)^*t*^, we introduce the following variable:

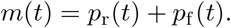

The variable *m*(*t*) represents the location in the direction of (1, 1)^*t*^ on the (*p*_f_, *p*_r_) plane. From the temporal change in *m*(*t*), we can observe the temporal change in the coordinates on the axis parallel to the iso-velocity lines.

We focused on the pressure dynamics on day 1 when crossing the b-Q. The initial period of the day was during stimulation, followed by a period without stimulation. It follows that the pressure transitions toward the black point, 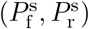, firstly, and then toward the white point, 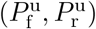 (Fig. 10(a)). Considering the behavior of *m*(*t*), the increase in *m*(*t*) under stimulation exceeds its decrease in the absence of stimulation when 0 *t <* 24 (Fig. 10(b)). Consequently, the initial state on Day 2 was positioned to the upper right of that on Day 1 (Fig. 10(a)). Next, we focus on the case where the crossing the b-Q is repeated. The increase in *m*(*t*) under stimulation gradually balanced its decrease in the absence of stimulation by repeatedly crossing the b-Q (Fig. 10(b)). Consequently, the displacement of the initial state to the upper right decreased and the deceleration converged (Fig. 6(b)).

**Fig. 10.**
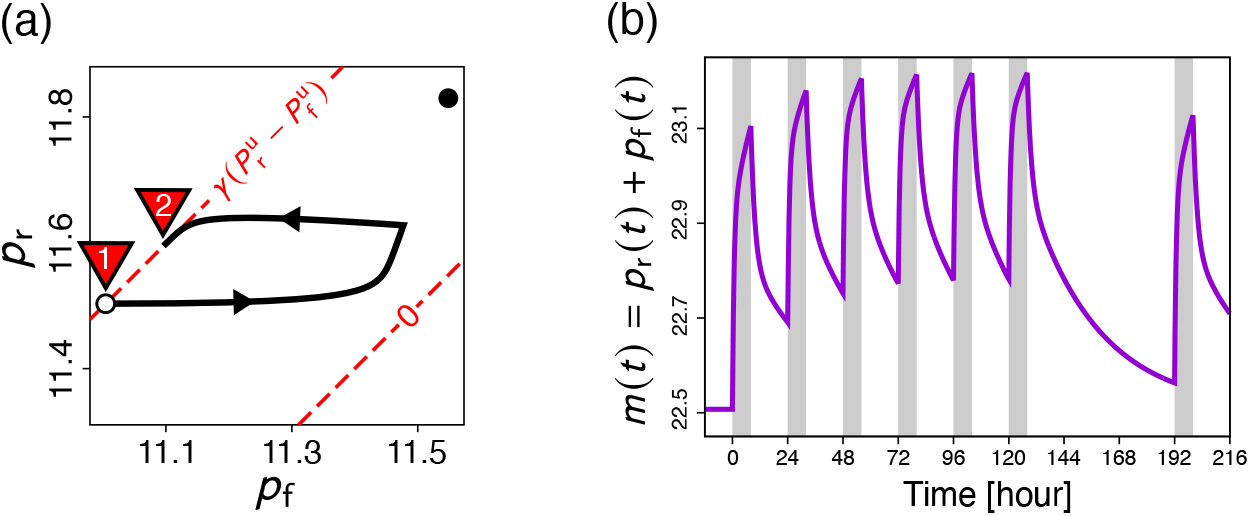
(a) The trajectory of day 1 on the (*p*_f_, *p*_r_) plane. The white circle represents 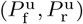, and the black circle represents 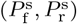. The red triangle with integer *n* points to the initial value at day *n*. (b) The temporal dynamics of *m*(*t*). Gray background represents the period under stimulation, *h*(*t*) = 1, and white background represents the period without stimulation, *h*(*t*) = 0

In summary, a shift in the initial state of each day is important for mitigation and recovery from deceleration. Additionally, we found that its mitigation caused a difference between the displacement to the axis parallel to the iso-velocity lines under stimulation and in its absence.

### 5.3 Velocity Consistency on the b-N

Here, we discuss the reason for the constant locomotion velocity of the b-N. The initial locomotion velocity on each day was almost the same as 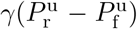 regardless of whether the bridge condition on the previous day was quinine or normal (Fig. 6(b)). In short, the initial states were sufficiently close to the straight line 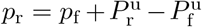, where the velocity was 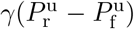 (Fig. 9). We now consider why the initial states are distributed in this manner.

The sol pressure (*p*_f_ (*t*), *p*_r_(*t*)) first approaches a straight line 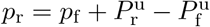 in the absence of stimulation (Fig. 7(b)). The eigenvalues of Eq. (10) are negative, and the absolute value of the eigenvalue corresponding to the eigenvector (1, −*r*_v_)^*t*^ is larger than that corresponding to the eigenvector (1, 1)^*t*^ due to the diffusion effect (Eq. (11) and (12)). Thus, the sol pressure decays faster toward (1, −*r*_v_)^*t*^ on the (*p*_f_, *p*_r_) plane. In addition, the sol pressure transitions along the straight line 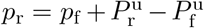 after approaching this line, because Eq. (10) has the eigenvector (1, 1)^*t*^ (Fig. 7(b)).

When crossing the b-Q, stimulation was absent during the last period of the day and occurred before this period. Therefore, the sol pressure returns to a straight line 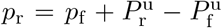 although it leaves the line for some time when crossing the b-Q (Fig. 9(a) and 10(a)). When crossing the b-N, the sol pressure transitions along the straight line 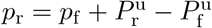 (Fig. 9(b)).

Consequently, the initial states are distributed along a straight line 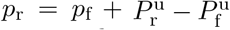 (Fig. 9). From this distribution, the sol pressure transitions along a straight line when crossing the b-N, independent of the bridge conditions of the previous day (Fig. 9(b)). Thus, the locomotion velocity remained constant (Fig. 6(b)).

## 6 Discussion

In the mathematical model proposed in this study, crawling migration (Eq. (1)) is regulated by the spatial difference in the protoplasmic pressure between the frontal and rear parts (Eq. (2) and (3)), and exhibits the two hallmarks defined in Table 1: habituation and spontaneous recovery.

In this section, we discuss the proposed model from a biological perspective. Deceleration mitigation as an adaptive behavior is realized by an initial state shift along the straight line 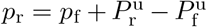. Chemical diffusion and spatial differences in chemical decomposition rates are important for adaptive behavior.

The diffusion effect contributed to the transition of the initial state. Owing to diffusion effects, *p*_r_ increases with increasing *p*_f_ under stimulation. This, in turn, causes the displacement of the sol pressure (*p*_f_, *p*_r_) parallel to the iso-velocity lines to increase (Fig. 7(a) and 9(a)). In the absence of stimulation, the sol pressure (*p*_f_, *p*_r_) returns to a straight line 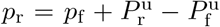 (Sec. 5.3). Based on these factors, an initial state shift along the straight line 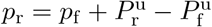 is realized.

The spatial difference in the chemical decomposition rate contributes to the connection between the initial state shift and mitigation of deceleration. The relationship between the decomposition rates under stimulation in the front and rear parts determines the slope of the iso-acceleration line Eq. (16). In our simulation, 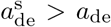 is true. Thus, the slope is greater than 1. However, the direction of the initial state shift is parallel to the iso-velocity line, whose slope is equal to 1 (Eq. (14)). Therefore, the direction of the initial state shift and the iso-velocity line intersect because 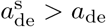 (Fig. 9(a)). Consequently, the deceleration was mitigated.

In summary, the diffusion effect between the front and rear parts is crucial for the transition of the initial state, and the difference in the chemical decomposition rate between the front and rear parts is important for reflecting the transition in the locomotion velocity. In short, the spatial scale of plasmodium played a key role in adaptive behavior in our model.

Some models exhibiting these two hallmarks (Table 1) have been proposed (Eckert et al. 2024; Stanley 1976). In a model based on the biochemical kinetics of cellular movement, two distinct chemical substances—activators and inhibitors—control the output (Eckert et al. 2024). In a model based on signal processing in neural circuits, the output is activated by the firing of other neurons and inhibited by a decline in the synaptic connection strength (Stanley 1976). In both models, the core mechanism controls the balance between two antagonistic factors: activation and inhibition of the response.

Although the biological realities represented by these models differ from those of our model, we considered whether there is any similarity in the core mechanism between our model and previously proposed models. In previous models, a common dynamic, which controls the balance between the inhibitory and activating effects on the response, was found.

In our model, the core mechanism lies in the relationship between the pressure difference of the protoplasm and the locomotion speed. In our simulation, the deceleration of locomotion speed under stimulation was decreased by repeated stimulation (Fig. 6(b)). Thus, we regarded deceleration during stimulation as a response to the stimulation. The variable *p*_f_ acts as an activator for deceleration under stimulation because a higher *p*_f_ decreases locomotion speed (Eq. (1)). In contrast, the variable *p*_r_ acts as an inhibitor of deceleration under stimulation because a higher *p*_r_ increases locomotion speed (Eq. (1)). In this sense of a dynamical system, our model has a mechanism similar to that of previous models.

However, the novel point worth noting in our model is that the two variables

of the activator and inhibitor are not different chemical components, but the same component in different places in plasmodium (Fig. 3). Spatial differences in identical biochemical processes can exhibit two hallmarks of habituation. In other words, our model suggested that the spatial inhomogeneity of a chemical plays a crucial role in the habituation of spatially extending cells, such as *P. polycephalum*.

We compared our model with that of Eckert et al. (2024) based on an incoherent feed-forward (IFF). They reproduced the two features of habituation that our model does (Table 1) using the IFF model. In their model, two factors, *I* and *M*, act on the response *R* (Fig. 11(a)). The activation effect under stimulation from *I* to *R* was constant for each stimulation (top of Figure 2C and 3B in Eckert et al. (2024)). However, the inhibitory effect under stimulation from *M* to *R* depends on the series of stimulations. For example, the inhibitory effect gradually increases with each stimulation (top of Figure 2C and 3B in Eckert et al. (2024)). This is because *M* increases with repeated stimulations. In contrast, the inhibition effect decreases when stimulation is withheld, owing to the decrease in *M* . In brief, the active effect on *R* was constant, but the inhibitory effect depended on the stimulation history. The IFF model realized two features of habituation that our model reproduced from the above properties.

**Fig. 11.**
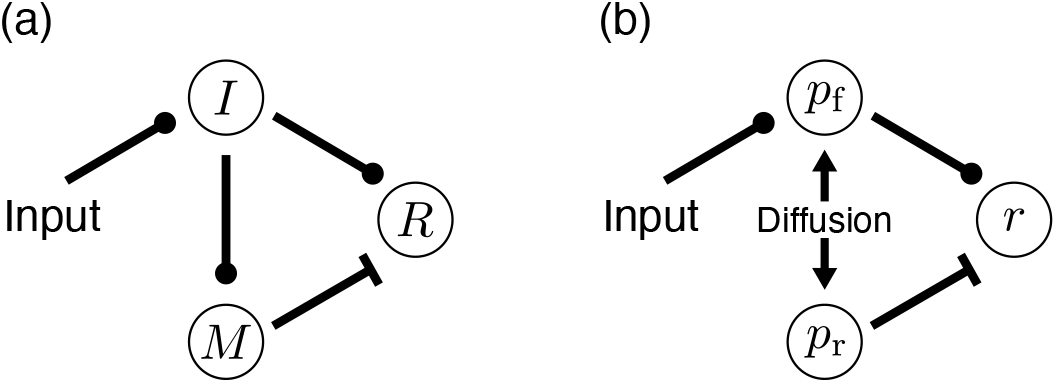
(a) The schematic diagram of IFF model in Eckert et al. (2024). (b) The schematic diagram of our model. *r* is equal to 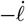. Line with a circle represents activation effect, line with a T-shaped end represents inhibition effect, arrow represents the generation effect of chemical and double-headed arrow represents diffusion effect

In our model, *p*_f_ and *p*_r_ affect the locomotion velocity 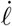. As previously mentioned, we defined deceleration under stimulation as the response to stimulation. To make it easier to compare the IFF model with ours, we regard an increase in 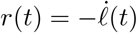 as the response. *p*_f_ is an activator of *r* and *p*_r_ is its inhibitor (Fig. 11(b)). *p*_f_ increased under stimulation. In addition, *p*_r_ increased under stimulation because of the diffusion effect (Fig. 7(a), 9(a), and 10(a)). For a more complex situation in which both the activator and inhibitor change, we analyzed the behavior of (*p*_f_ (*t*), *p*_r_(*t*)) in the phase plane. The initial state shift was important for the decline and recovery of the response (Sec. 5.2 and 5.3). In our model, stimulation history was reflected in the initial state shift. To summarize, a common feature of the IFF model and our model is that they include a structure that stores the stimulation history. In the IFF model, variable *M* (*t*) plays the role of memory. In our model, this is the initial state transition in the phase plane.

In the behavioral definition of habituation, there are ten hallmarks (Rankin et al. 2009). Some models have already been proposed to reproduce three or more hallmarks, but these models are more complicated than our model, as they involve more variables and incorporate nonlinearity in ODEs (Eckert et al. 2024; Smart et al. 2024), whereas our model essentially comprises two linear ODEs with switching parameters (Eq. (2) and (3)). For example, Eckert et al. (2024) reproduced other hallmarks by concatenating two IFF models. Our model was constructed in a bottom-up approach and it has the potential to be improved by adding new variables and/or nonlinearities, thereby reproducing three or more hallmarks.

In this study, we analyzed the behavior of (*p*_f_ (*t*), *p*_r_(*t*)) on the phase plane to elucidate the fundamental structure of the two hallmarks: habituation and spontaneous recovery (Table 1). The analysis revealed that the initial state transition on an isovelocity line is important for two hallmarks (Sec. 5.2 and 5.3). Our model provides an understanding of these two hallmarks from the perspective of a dynamic system. Our future research direction will be to analyze other hallmarks of habituation from the same perspective. This approach will bring new progress toward a comprehensive understanding of habituation mechanisms among a wide range of species.

## A Derivation of the Diffusion Term

We derived the diffusion term for different solution volumes. We consider adjacent solutions 1 and 2 (Fig. 12). The volume of solution *i* was defined as *V*_*i*_ [m^3^]. The chemical concentration of solution *i* is defined as *c*_*i*_ [mol*/*m^3^]. *J*_*i*_ represents the flux [mol*/*m^2^*·*s]. We derived the diffusion term when *V*_1_ ≠ *V*_2_. In this study, we assumed that the flux from both ends is zero; that is, *J*_0_ = *J*_2_ = 0. From Fick’s first law, *J*_1_ can be expressed as follows:

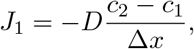

where *D* represents the diffusion coefficient [m^2^*/*s], and Δ*x* is the distance between solutions 1 and 2 [m]. Note that the flux *J*_*i*_ is positive to the right. The inflow and outflow of solute particles per unit time in solution 1 are *S*(*J*_0_ *J*_1_) and where *S* denotes the particle passage. Thus, the following equation is obtained:

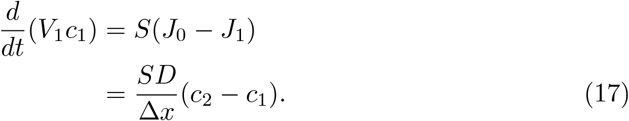

**Fig. 12.**
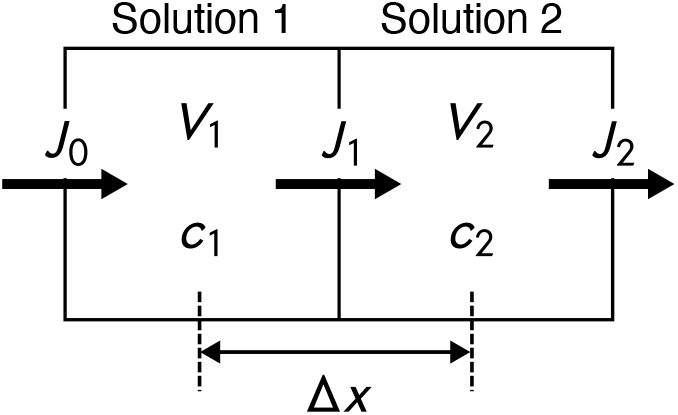
The problem setting of the diffusion between the different volumes of solutions. Arrow represents the flux

From Eq. (17), the differential equation for *c*_1_ is derived as

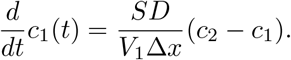

Similarly, the differential equation for *c*_2_ is derived as follows:

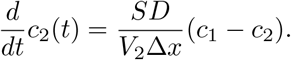

We now define *d* and *r*_v_ as follows:

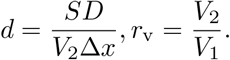

Ultimately, we obtain the differential equation with the diffusion term between different volumes of the solutions as follows:

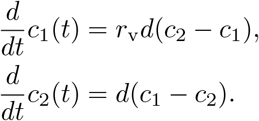

## Acknowledgements

K. N. was supported by the Doctoral Program for World-Leading Innovative & Smart Education (WISE program) from the Ministry of Education, Culture, Sports, Science, and Technology (MEXT) at Kyushu University. This work was supported by the Cooperative Research Program of the “Network Joint Research Center for Materials and Devices” of MEXT, Japan.

## Statements and Declarations

## Authors’ Contribution

A.T. proposed the fundamental ideas for the mathematical model, while K.N. performed the formulation, simulation, and analysis under A.T.’s supervision. T.N. and

Y.N. examined the biological validity of the model. K.N. drafted the manuscript, which was revised by T.N. and Y.N. Throughout the entire research process, all authors collaborated and advanced the work through discussions under A.T.’s leadership.

## Data Availability

The experimental data used in this study were originally generated by Boisseau et al. (2016) and are publicly available at https://doi.org/10.5061/dryad.51j89.

